## Supplementary figures for "Systematic evaluation of intratumoral and peripheral BCR repertoires in three cancers"

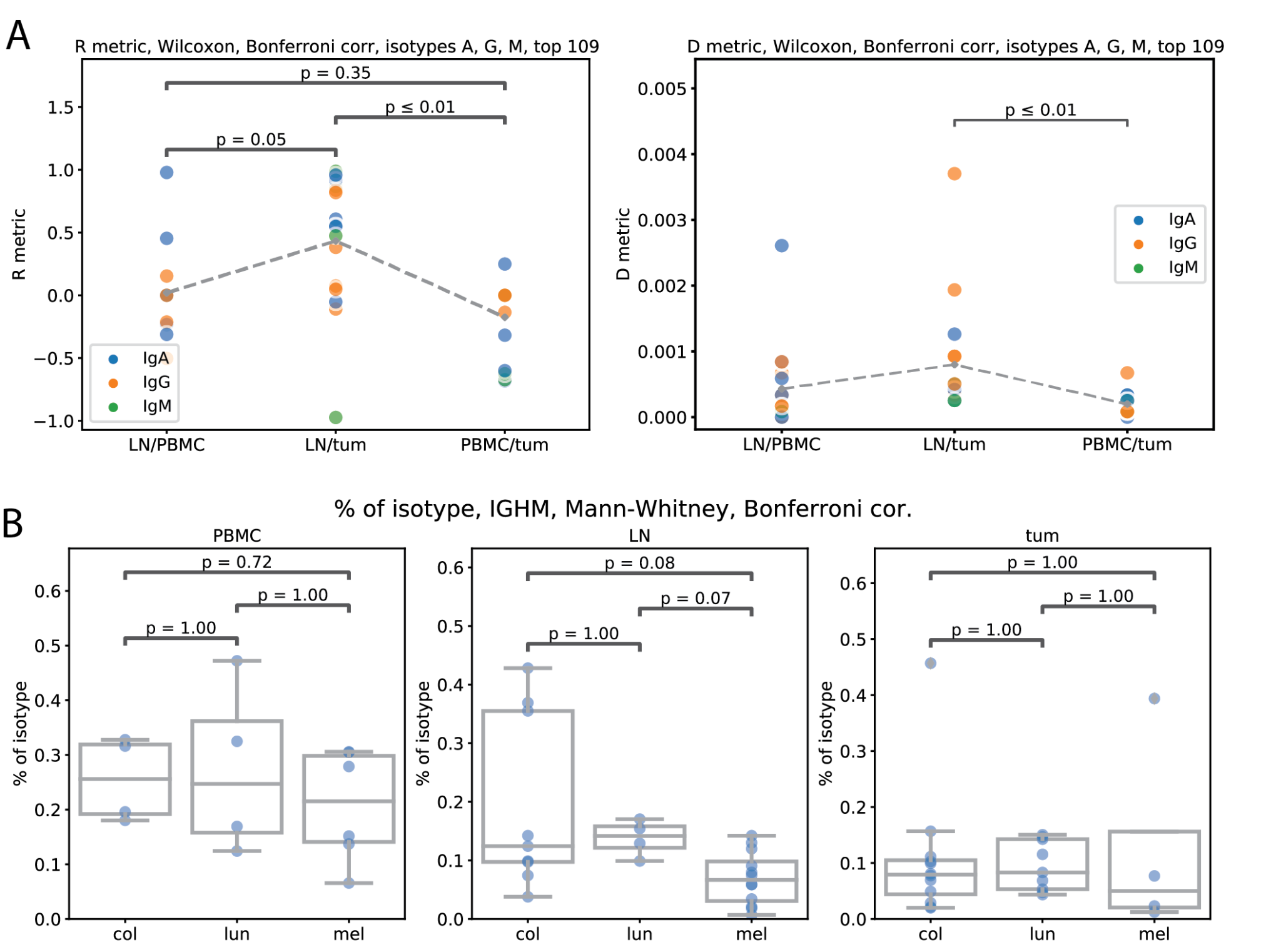

#### **Figure S1 Repertoire similarity by R metric, repertoire overlap by D metric (A), isotype composition by tissue type and cancer (B).**

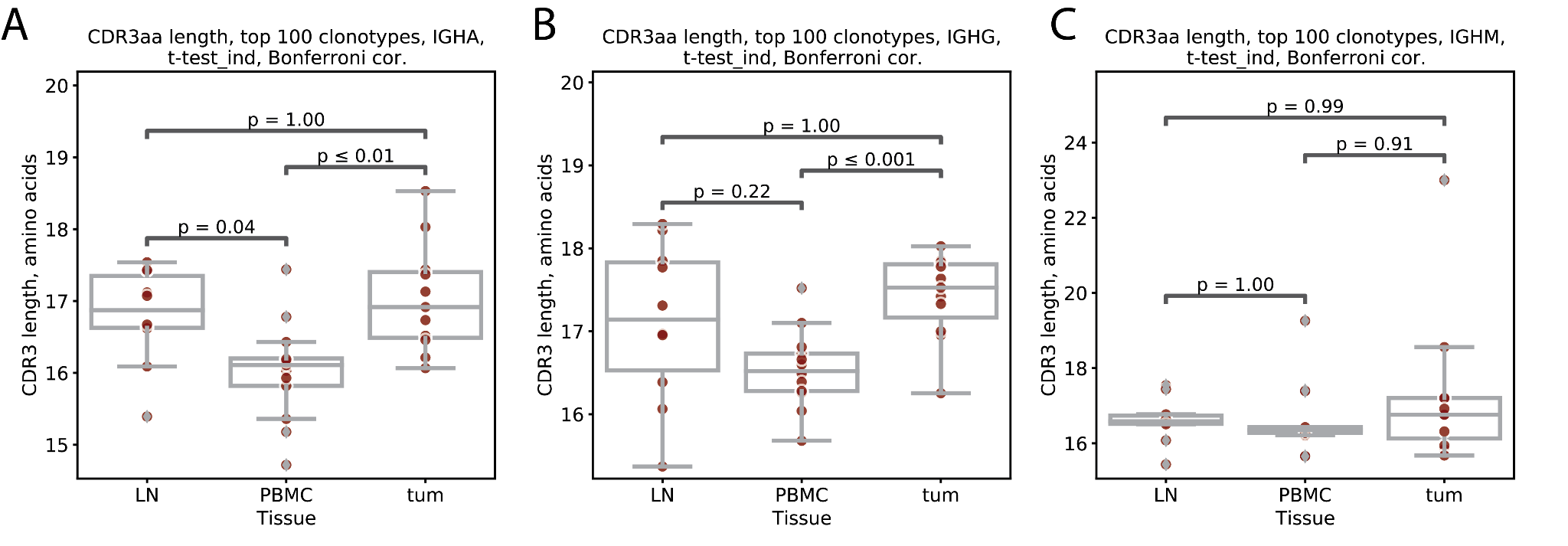

#### **Figure S2 Mean CDR3 length for IGHA, IGHG and IGHM repertoires from lymph node, PBMC and tumor.**

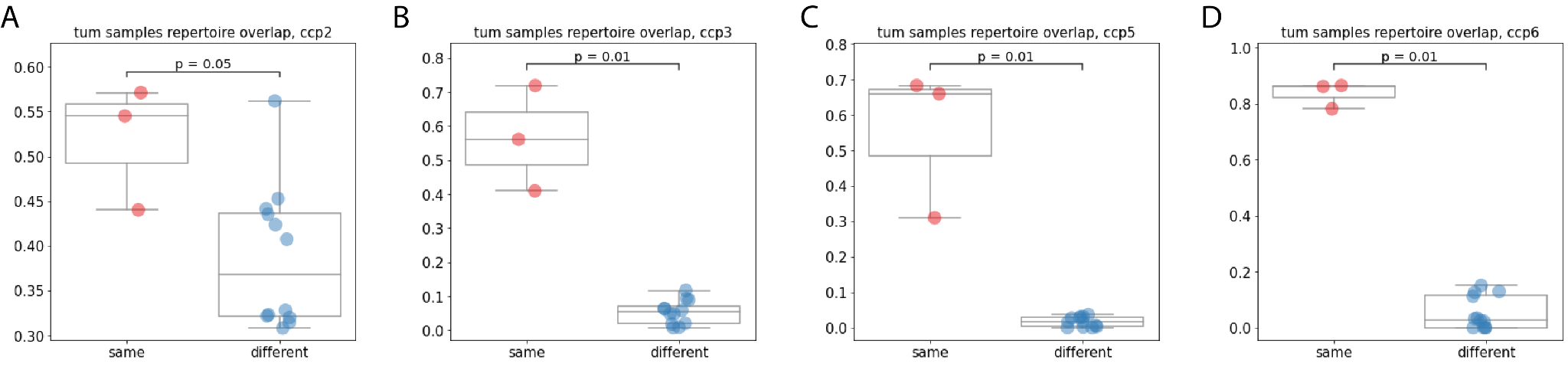

#### **Figure S3 BCR repertoire overlap for separate fragments of tumor (different), and from cell suspension level replicates from the same fragment (same), patients ccp2 (A), ccp3 (B), ccp5 (C), ccp6 (D).**

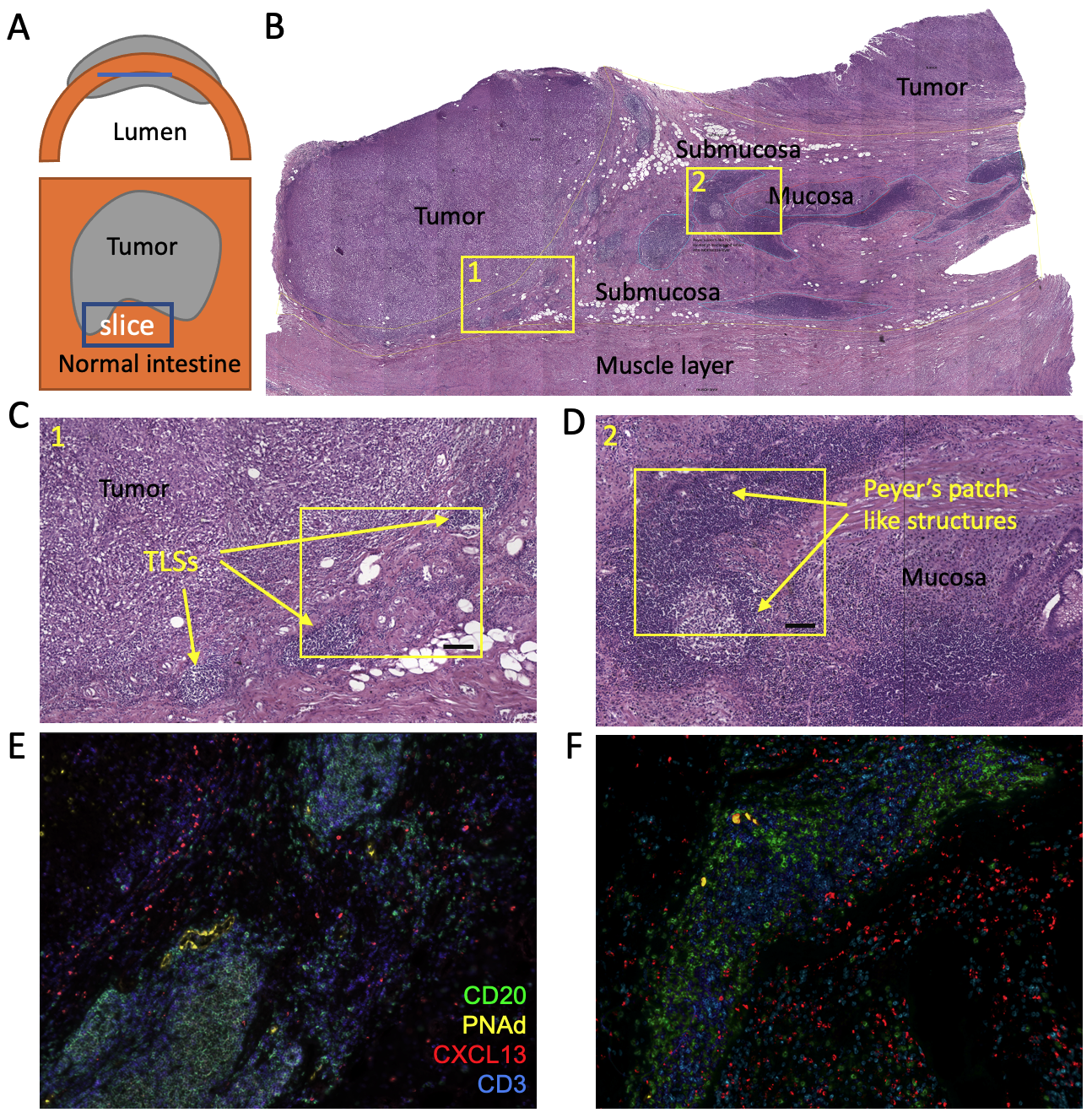

#### **Figure S4. Histochemical analysis of lymphoid infiltration in ccp2 sample. A. Schematic representation of slice orientation relative to the intestine wall and tumor. B. Tissue section stained with hematoxylin and eosin (H&E). Different layers of the intestine wall and tumor are marked. Regions 1 and 2 are magnified on the panels C and D, respectively. C, D. H&E histology of the peritumoral region and mucosal/submucosal layers. Lymphoid structures in these regions were classified as TLS and Peyer’s patches, respectively and are marked with arrows. Regions on C and D correspond to the areas that are shown on fluorescent images of parallel slices in E and F, respectively. CD20 fluorescence is shown in green, CD3 in blue, CXCL13 in red and PNAd in yellow. Scale bar is shown in black and corresponds to 100 µm.**

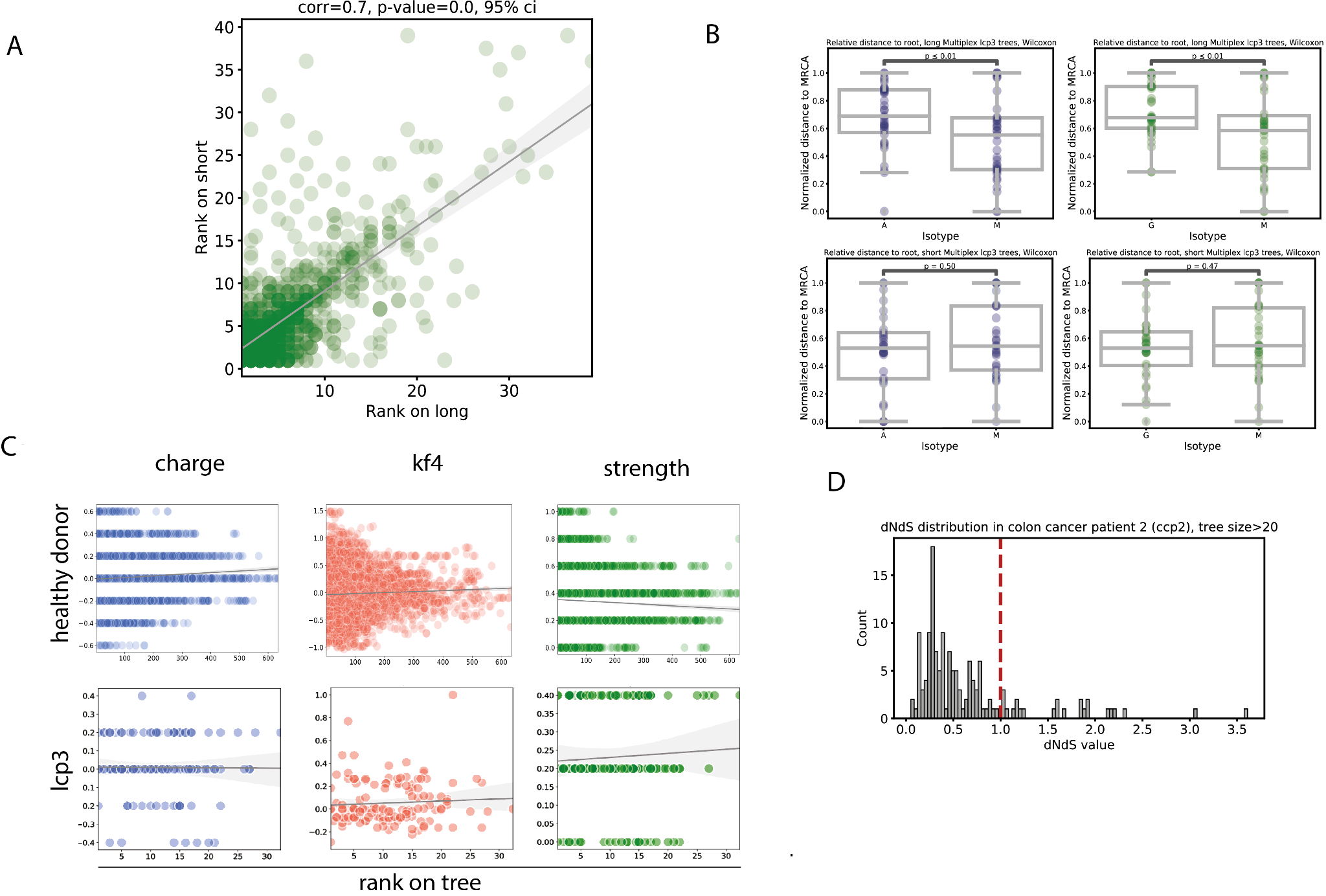

#### **Fig.S5 Short (CDR-H3-based) versus full-length IG trees phylogeny analysis**

**A** - distance to MRCA comparison for same clonal groups on short and long trees; **B** - distance to MRCA comparison for IgM, IgA and IgG-dominated clonal groups on short and long trees; **C** - distribution of CDR-H3 amino acid properties (charge, kf4 and strength) along short and long trees in healthy donor (top row) and cancer patients (bottom row); **D** - dN/dS values for trees os size > 20

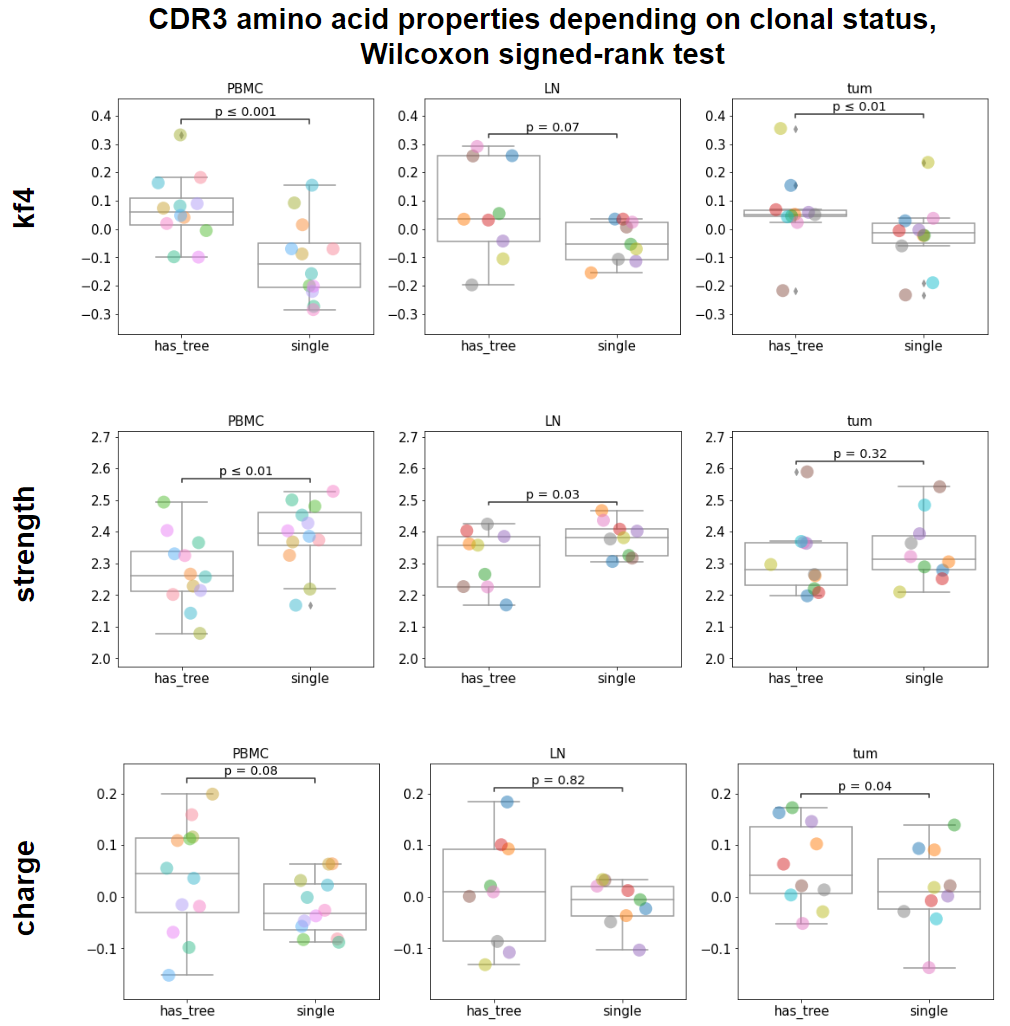

#### **Fig.S6 aa properties depending on clonal status**

Kf4, strength and charge for central 5 aa of CDR3 region in repertoires from PBMC, LN and tumor, for CDR3 clonotypes that belong to a clonal group (has_tree) and those that don’t belong to any clonal group (single).
